# Strong epistasis constrains the evolution of strict substrate specificity in apicomplexan lactate dehydrogenases

**DOI:** 10.1101/2023.06.21.545977

**Authors:** Jacob D. Wirth, Jeffrey I. Boucher, Changhan Xu, Scott Classen, Douglas L. Theobald

## Abstract

The homologous enzymes lactate and malate dehydrogenase (L/MDH) are structurally similar but are specific for different substrates. LDH vs MDH specificity is canonically governed by the identity of a single “specificity residue” at position 102. However, LDH function has convergently evolved from a specific MDH at least four times, and the catalytic role of residue 102 is not conserved between different phyla. The apicomplexa are a phylum of obligate, intracellular eukaryotic parasites responsible for wide-spread disease such as *Plasmodium falciparum* (malaria), *Cryptosporidium parvum* (cryptosporidiosis), *Toxoplasma gondii* (toxoplasmosis), and *Eimeria maxima* (eimeriosis). The apicomplexan LDH evolved via a five-residue insertion that produced a novel specificity residue, W107f. The commonly accepted mechanism of LDH specificity involves charge balance and steric occlusion, but our data shows that the general mechanism of apicomplexan LDHs does not use W107f as a steric block. Only *Plasmodium* LDHs evolved substantial steric specificity, making them exceptional among Apicomplexa. Strong protein epistasis constrained this evolution, making it difficult to revert to ancestral phenotypes. Here, we use ancestral sequence reconstruction (ASR), steady-state kinetics, and x-ray crystallography to characterize apicomplexan LDHs which challenge current assumptions about the evolution of L/MDH activity. We demonstrate the unique specificity of *Plasmodium* LDHs and identify the active site residues controlling their substrate recognition. The extraordinarily high specificity of *Plasmodium* LDHs presents difficulties for small-molecule inhibitor development, and successful drugs against Plasmodium LDH may not be efficacious against other Apicomplexa LDHs and their diseases.

## Introduction

Protein epistasis is a widely occurring phenomenon where the effect of a mutation is dependent on its genetic context, in this case defined as the presence of other residues^1-2^. Epistasis can be positive (permissive) or negative (restrictive) and in some cases cause a mutation’s effect to change from beneficial to detrimental (or vice versa)^3^. Nonspecific epistasis may describe a many-to-many relationship that involves a large amount of a protein’s sequence while more specific epistasis could involve the presence of a single permissive residue allowing for the mutation of another^2^. The direct effect of an epistatic relationship is witnessed when mutating the equivalent position in two closely related proteins has significantly different outcomes.

Lactate and malate dehydrogenases (LDHs and MDHs) are homologous metabolic enzymes that share a protein fold^4^ and a common catalytic mechanism (Figure 1)^5-8^. LDH interconverts pyruvate and lactate in anaerobic glycolysis. In the presence of oxygen, metabolites are instead driven through the citric acid cycle where MDH interconverts oxaloacetate and malate. Their roles in central metabolism make them essential and found throughout all three domains of life^9^. The shared catalytic mechanism and ordered bi-bimolecular kinetic scheme stem from the family’s highly conserved active site and similar substrates (Figure 1). The enzymatic reduction of substrate proceeds as follows: 1) NADH binds free enzyme, 2) R171 coordinates the substrate’s carboxylic acid group, 3) a loop closes over the active site, binding the substrate, 4) hydride transfer occurs, 5) the active site loop opens, ejecting the reduced product, and 6) the enzyme regenerates by releasing NAD^+^. D168 is required to activate the catalytic H195, which is directly involved in the proton transfer during catalysis. Research has shown that the movement of the active site loop, known as the specificity loop, is the reaction’s primary rate-limiting step^10-11^.

**Figure 1.**
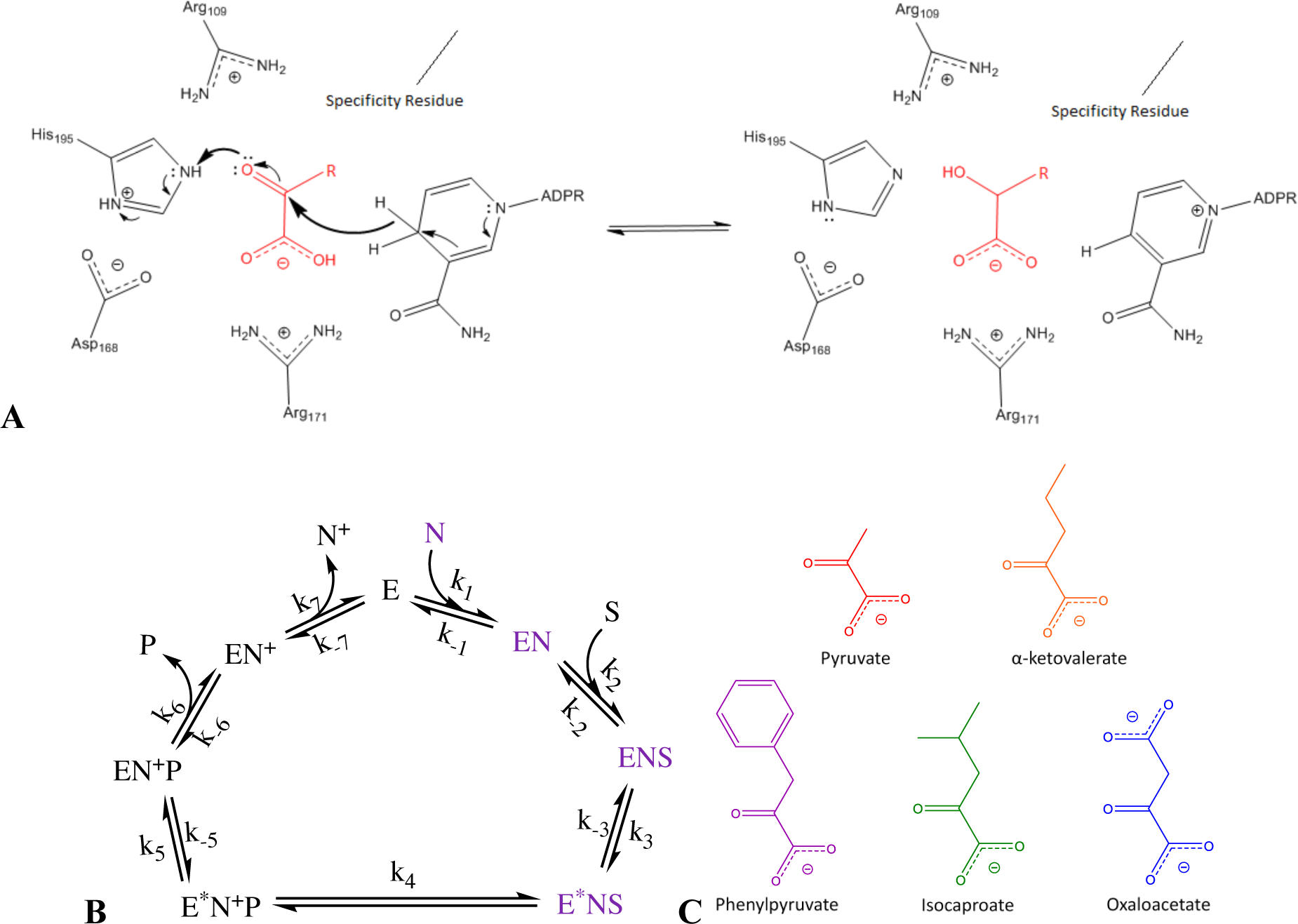
Mechanism of loop closure and catalysis. **(A)** and **(B)** are the molecular and kinetic mechanism of substrate turnover for LDHs and MDHs as described in the text. Specificity residues vary. **(C)** Alternate substrates can be used to study LDH and MDH specificity.

The loop is so-named because it plays an important role in substrate recognition. LDHs and MDHs have strict specificity for their respective substrates, though occasionally substrate analogs (Figure 1c) are turned over with modest activities^10, 12-14^. Residue 102 resides on the loop and is commonly referred to as the ‘specificity residue’ because it differentiates between the functional groups of different α-ketoacids^11-12, 15^. MDHs have R102 which can form a salt bridge with the γ-carboxylate group of its native substrates, oxaloacetate and malate. The most common form of LDH, like those in bacteria and metazoa, contain Q102 which lacks arginine’s formal positive charge and allows for stable binding of pyruvate or lactate^12^.

The enzyme family has often been a model for functional evolution studies with many descriptions, solved protein structures, and documented functional changes^16-18^. No two types of LDH naturally use the same specificity residue(s) for activity, suggesting that each group evolved a different form of substrate discrimination. In some cases it is possible to swap specificities between LDH and MDH^12, 19^, while in others it is only marginally effective^20^ or produces completely inactive enzymes^21^. As LDHs evolved from MDHs multiple independent times^14, 18, 22-23^, these specificity studies support the idea that epistasis has strongly constrained the evolution of LDH activity which in turn could explain why each type of LDH required very different mutations to develop pyruvate specificity.

One independent family of LDHs belongs to the intracellular eukaryotic parasites Apicomplexa, human pathogens that cause many diseases including malaria, cryptosporidiosis, babesiosis, and toxoplasmosis. The organisms responsible for malaria belong to the *Plasmodium* genus with the most fatal form of malaria due to *Plasmodium falciparum*. *P. falciparum* proceeds through a complex life-cycle in two different hosts where *P. falciparum* sporozoites infect humans *via* a mosquito bite. LDH in *Plasmodium* is transcribed at all stages in blood^24^ and has been measured in the micromolar range^25^ indicating the enzyme likely performs a critical role. While in host blood, *P. falciparum* respires anaerobically to regenerate NAD^+^ due to the conditions of the human red blood cell^26-27^. It is likely that LDH is the only means of NAD^+^ regeneration during this time, making the enzyme essential for the pathogen’s survival.

LDHs in Apicomplexa evolved nearly one billion years ago^28^ from a horizontal gene transfer of an ancient α-proteobacterial malate dehydrogenase, independently of canonical metazoan and bacterial LDHs^18, 29-30^. The gene transfer occurred before Apicomplexa started to speciate and so all apicomplexan LDH genes, save one genus, come from one convergent event and have similar characteristics^22^. For the purpose of this study, LDHs from *Cryptosporidium* will not be addressed as they have evolved separately^22^. Apicomplexan LDHs did not evolve by mutating a glutamine at position 102, despite sharing the human LDH catalytic mechanism^10^. Rather, they evolved from an ancestral MDH through a unique five amino acid insertion in the substrate specificity loop that switches substrate specificity from malate/oxaloacetate to lactate/pyruvate^23^. The insertion lengthens the loop but otherwise the structures of apicomplexan and metazoan LDHs are highly similar^31^.

A canonical human-like LDH structure with the substrate specificity loop in the closed conformation has the Q102 specificity residue contacting substrate^32^. In contrast, the *P. falciparum* LDH (LDH_PLFA) crystal structure shows W107f contacting a substrate analog in the closed conformation, which indicates that W107f is the ‘specificity residue’ Apicomplexa evolved instead of something at position 102^33^. Point mutations and deletions within the substrate specificity loop have demonstrated that the essential W107f is the only crucial residue when determining enzyme activity and substrate recognition, while mutating K102 has negligible effects^23, 34^.

As stated above, pyruvate specificity in canonical LDHs or MDHs is decided by position 102 on an active site loop (Q102 or R102, respectively)^12^. We can rationalize W107f as a specificity determining residue as it satisfies the charge balance requirements of pyruvate turnover and can sterically occlude oxaloacetate from the closed conformation of the active site (Figure 2)^33^. Apicomplexan LDHs lack any activity towards oxaloacetate despite there being an available positive charge at K102, leading many to question why the enzymes do not also function as MDHs^10, 25, 35-38^. More puzzling, our group was able to make a bifunctional apicomplexan LDH through the mutation K102R, calling in to question exactly how the loop-insertion is used by the enzyme to determine specificity^23^.

**Figure 2.**
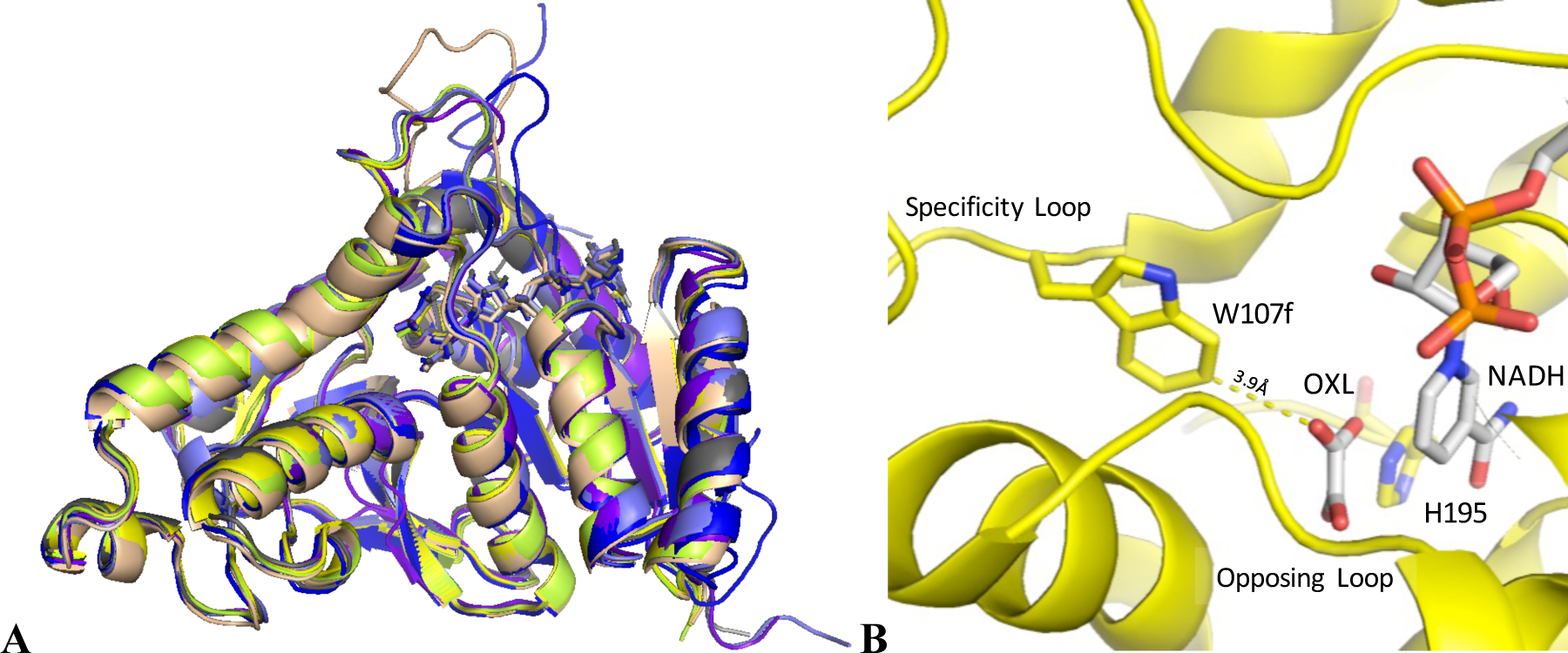
Apicomplexan LDH protein structure. **(A)** Six LDH structures from the Apicomplexa phylum exhibit highly similar global monomer structures: one *Eimeria*, two *Toxoplasma,* and three *Plasmodium* (PDBID: 6CT6, 1PZH, 1SOW, 1T2D, 2A92, and 1OC4, respectively). **(B)** The specificity loop with W107f is labeled, as well as a visible loop seen orientated across the active site. The pyruvate analog oxalate and NADH are shown as white sticks.

Noted previously, LDH_PLFA is unable to turnover large substrate analogs^10^, a behavior that we show is atypical for Apicomplexa. Instead, LDH_PLFA has exceptionally high specificity when compared to closely related LDHs from other Apicomplexa such as *Babesia bovis* (LDH_BABO), *Toxoplasma gondii* (LDH_TOGO), and *Eimeria maxima* (LDH_EIMA). LDH_PLFA has frequently been a target for small-molecule drug therapies due to its essential role during the parasite’s life cycle and active site features distinct from human LDH^33, 35, 39-45^. Any difficulties developing small-molecule drug therapies meant to target *Plasmodium* LDHs should be unsurprising if the organisms evolved an important but unexplored or uncharacterized mechanism of substrate specificity. Could that specificity be easily relaxed or changed? Could other LDHs easily evolve higher specificity?

Apicomplexa LDHs provide a good model system for investigating several questions in molecular evolution, such as the number of mutations required to change a protein’s phenotype, how enzymes evolve strong specificities, and to what degree epistasis affects the course of functional evolution. Here we use ancestral sequence reconstruction, steady-state enzyme kinetics, and x-ray crystallography to characterize LDHs from several Apicomplexa, including modern proteins, ancestral constructs, and evolutionary intermediates. We identify what attributes are unique in *Plasmodium* LDH active sites and their contribution to substrate specificity. We also demonstrate that the specific order of mutation has a large impact when maintaining an enzyme’s activity as it evolves new specificity.

## Results

### Substrate Specificity in Apicomplexan LDHs

The closely related extant LDH_PLFA and LDH_EIMA were assayed for their steady-state activities towards several substrates. Both enzymes have high activity using the native substrate pyruvate yet LDH_PLFA is significantly more specific. Compared to other LDHs from Apicomplexa, LDH_PLFA has a k_cat_/K_M_ higher for pyruvate and much lower when using other substrates (Figure 3, Supp. Table 1). LDH_EIMA greatly prefers pyruvate over oxaloacetate but has a more relaxed specificity overall, making it more like close homologs^36^. To generalize the behavior of all apicomplexan LDHs, our group reconstructed the sequence of their last common ancestor as described in the methods (Figure 4)^23^. LDH_EIMA and the reconstructed ancestor (AncLDH) have qualitatively identical phenotypes: enzyme activity decreases as the substrate R-group loses planarity and increases in size. In general, most apicomplexan LDHs are not as sterically specific towards their substrates as LDH_PLFA.

**Figure 3.**
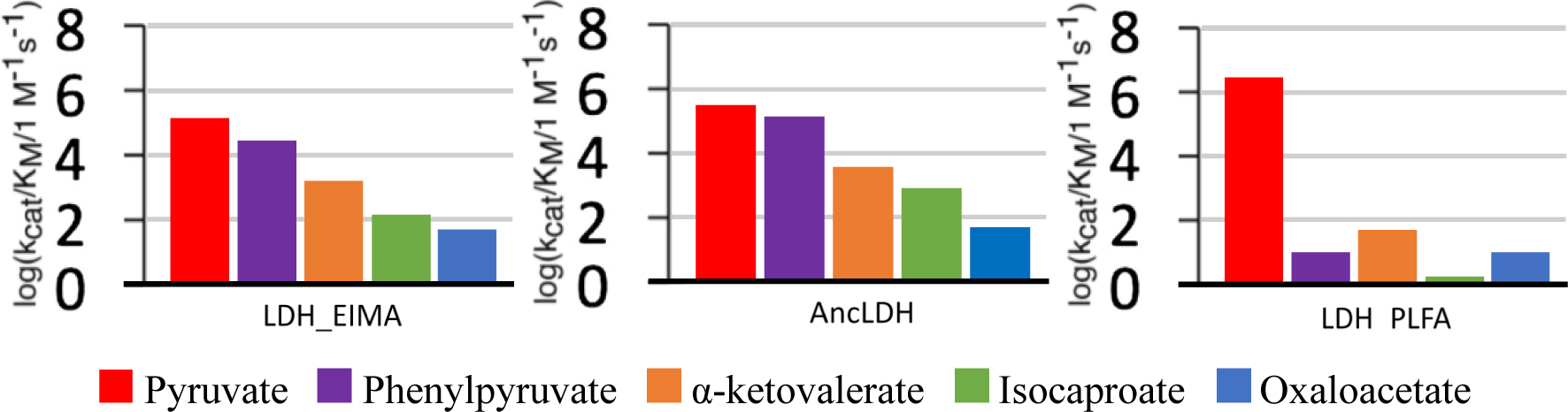
Apicomplexan LDH substrate specificity. Values are log(kcat/KM) towards α-ketoacid substrates in units of M^-1^*s^-1^. Each bar is colored by substrate. AncLDH is the reconstructed last common ancestor of all apicomplexan LDHs.

**Figure 4.**
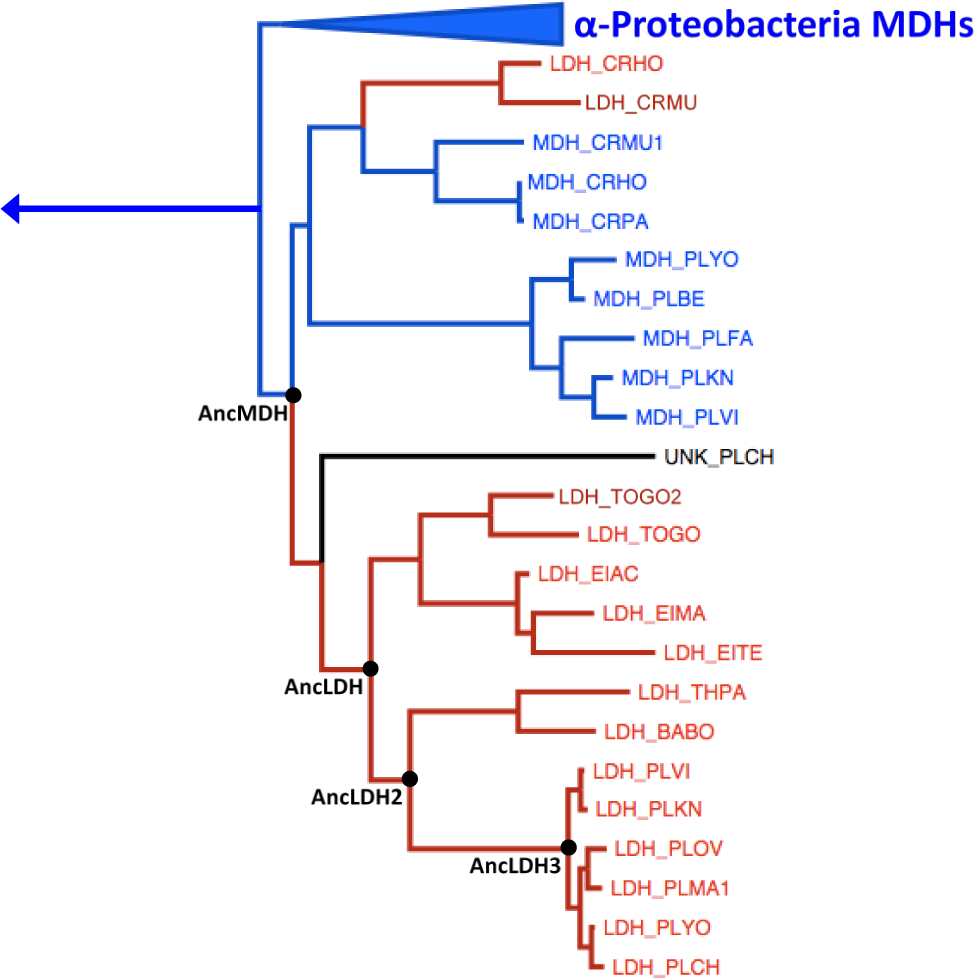
The apicomplexan L/MDH clade. MDHs are labeled blue and LDHs are red. The relevant reconstructed nodes are labeled in black.

LDH_PLFA’s inability to turnover phenylpyruvate was recorded previously^10^ but the mechanism behind such robust specificity was not explored. The large planar R-group is uncharged, allowing for use as a suitable substrate mimic to probe steric vs charge sensitivity in LDHs. Two additional LDH ancestors were reconstructed (AncLDH2 and AncLDH3, Figure 4) to follow the lineage towards *Plasmodium* LDHs and determine when the organisms gained the increased specificity. Along with another extant enzyme, LDH_BABO, the ancestral proteins were assayed for their pyruvate, oxaloacetate, and phenylpyruvate activities (Figure 5, Supp. Table 2). Apicomplexan MDHs are very specific for oxaloacetate, meaning the five-residue insertion is required for pyruvate turnover. It is apparent that the insertion also confers high phenylpyruvate activity which implies loss of specificity for oxaloacetate is due to charge and not size. Only LDH_PLFA and AncLDH3 were unable to use phenylpyruvate as a substrate which makes their steric discrimination a recent development in Apicomplexa.

**Figure 5.**
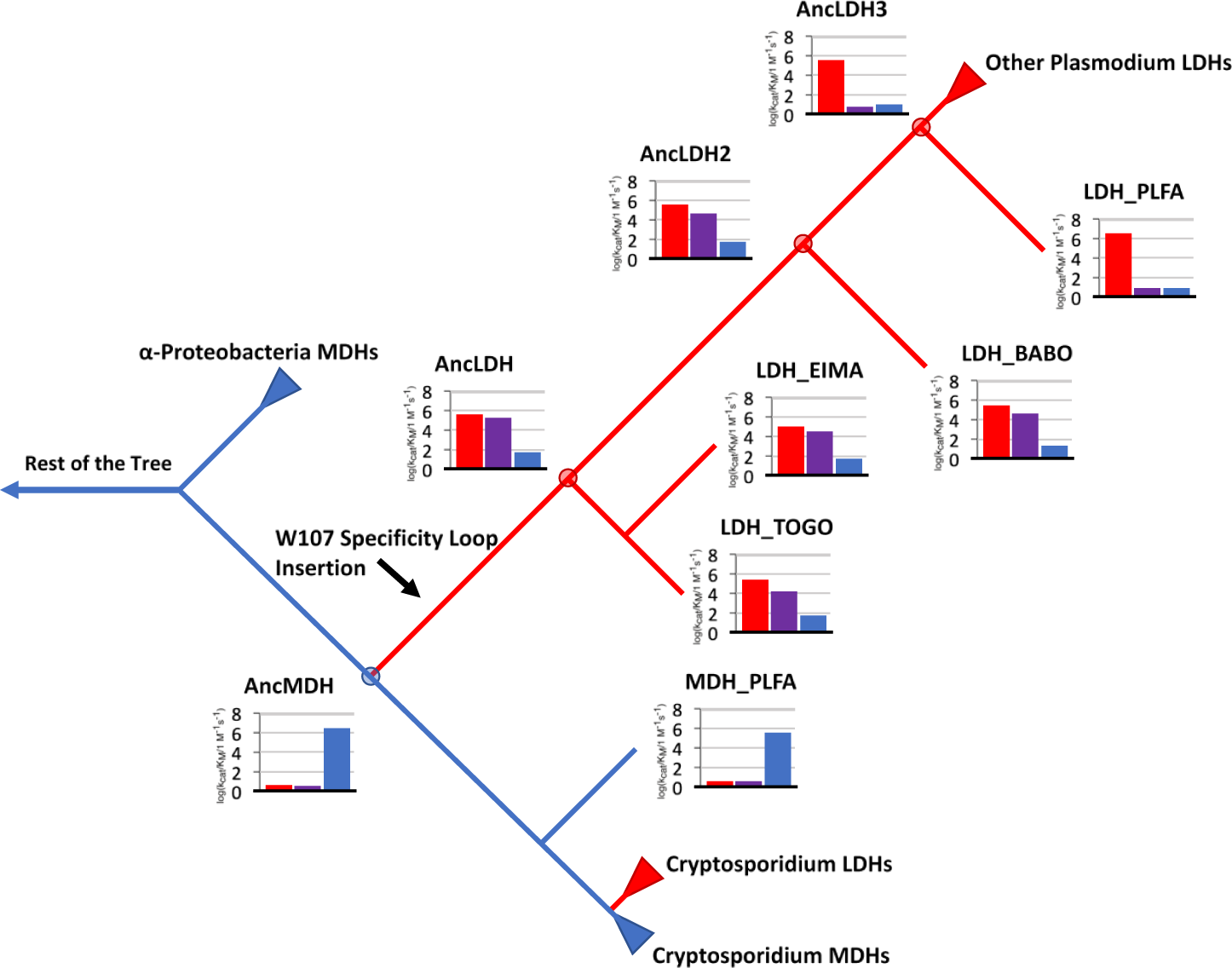
Evolution of Apicomplexa L/MDH Activity. The modern apicomplexan LDHs, MDHs, and ancestral reconstructions are plotted according to their phylogenetic relationship. The *Plasmodium* LDH and the *Plasmodium* ancestor have no phenylpyruvate activity. The coloring is consistent with Figures 3 and 4. Branch lengths are not to scale.

### Novel Active Site Residues in Plasmodium LDH

Comparing apicomplexan LDH crystal structures reveals a potential explanation for *Plasmodium’s* increased steric specificity. A *Toxoplasma* LDH crystal structure was available as an extant comparison and by using that, along with our solving of the LDH_EIMA structure (PDBID: 6CT6, Supp. Table 3), we were able to identify active site features unique to *Plasmodium* LDHs (Figure 6). Within the crystal structures, high steric specificity correlates with the identity of an active site loop opposite the specificity loop, between the α-G2 and α-G3 helices (the ‘opposing loop’, Figure 2b and 6b)^23, 33, 37-38, 46-47^. The opposing loop in most apicomplexan LDHs has a GQG motif (or GNG for LDH_BABO) but in *Plasmodium* LDHs there is a single alanine in the same three-dimensional space. Position 236 and 246 identity also appears mutated in the *Plasmodium* genus: G236A and A246P. The alanine and proline are both within 4 Å of pyruvate and as such are likely involved with substrate binding interactions (Figure 6c).

**Figure 6.**
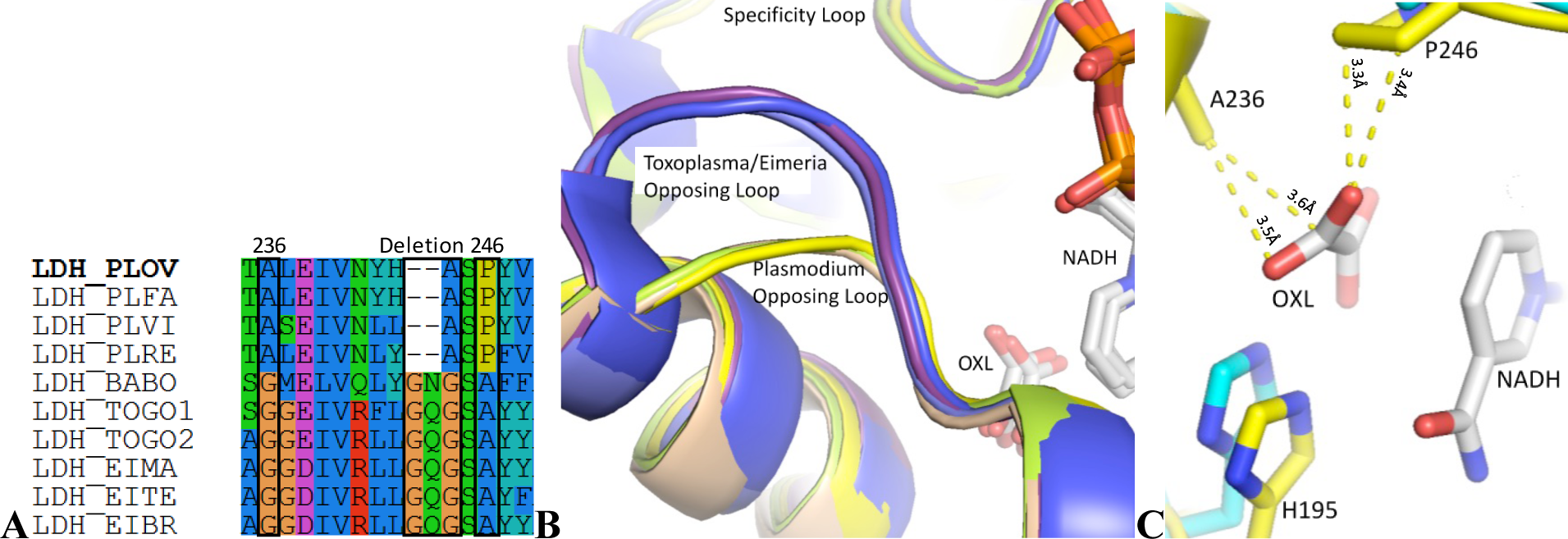
Comparison of apicomplexan LDH active sites. **(A)** Sites 236, 246, and opposing loop residues are generally slow-mutating positions (squared in black). Only *Plasmodium* LDHs have altered sequence identity when compared to other Apicomplexa. **(B)** *Plasmodium* LDHs have a novel deletion in an active site loop (structures from Figure 2). **(C)** Two residues are otherwise highly conserved in Apicomplexa. The unique *Plasmodium* residues A236 and P246 are within 4 Å of pyruvate in the *P. falciparum* LDH structure (Yellow; PDBID: 1T2D).

### Effects of Single Mutations

Using site-directed mutagenesis, we tested each position of interest for its contribution to steric specificity as well as all possible combinations. Trying to grant LDH_PLFA ancestral levels of promiscuity by mutating the active site residues abolishes enzyme activity to a non-measurable range. Epistasis in protein evolution is well-documented and an extant enzyme resisting reversion ‘backward’ through time to an ancestral state is unsurprising^1-2^. The relevant mutations in our ancestral constructs are less detrimental.

AncLDH2 has effectively no preference between pyruvate and phenylpyruvate (Figure 5 and 7). There is a decrease of five orders of magnitude in ability to turnover large substrates between AncLDH2 and AncLDH3 with almost no change in pyruvate rates. Each active site mutation incorporates into AncLDH2 with a minimal effect on pyruvate activity (Figure 7a). Phenylpyruvate turnover is reduced in AncLDH2_G236A by more three than orders of magnitude (Figure 7b). While this is a considerable drop, phenylpyruvate turnover is still ∼10-fold higher than in wild-type AncLDH3, so other residues must also be involved. A246P and the loop deletion have hardly any effect on the AncLDH2 construct. AncLDH3 is resistant to reversion but not as much as its modern counterpart. Pyruvate activity is lowered by a factor of 16 or 130 by mutating positions 236 or 246, respectively, with no accompanying increase in phenylpyruvate recognition.

**Figure 7.**
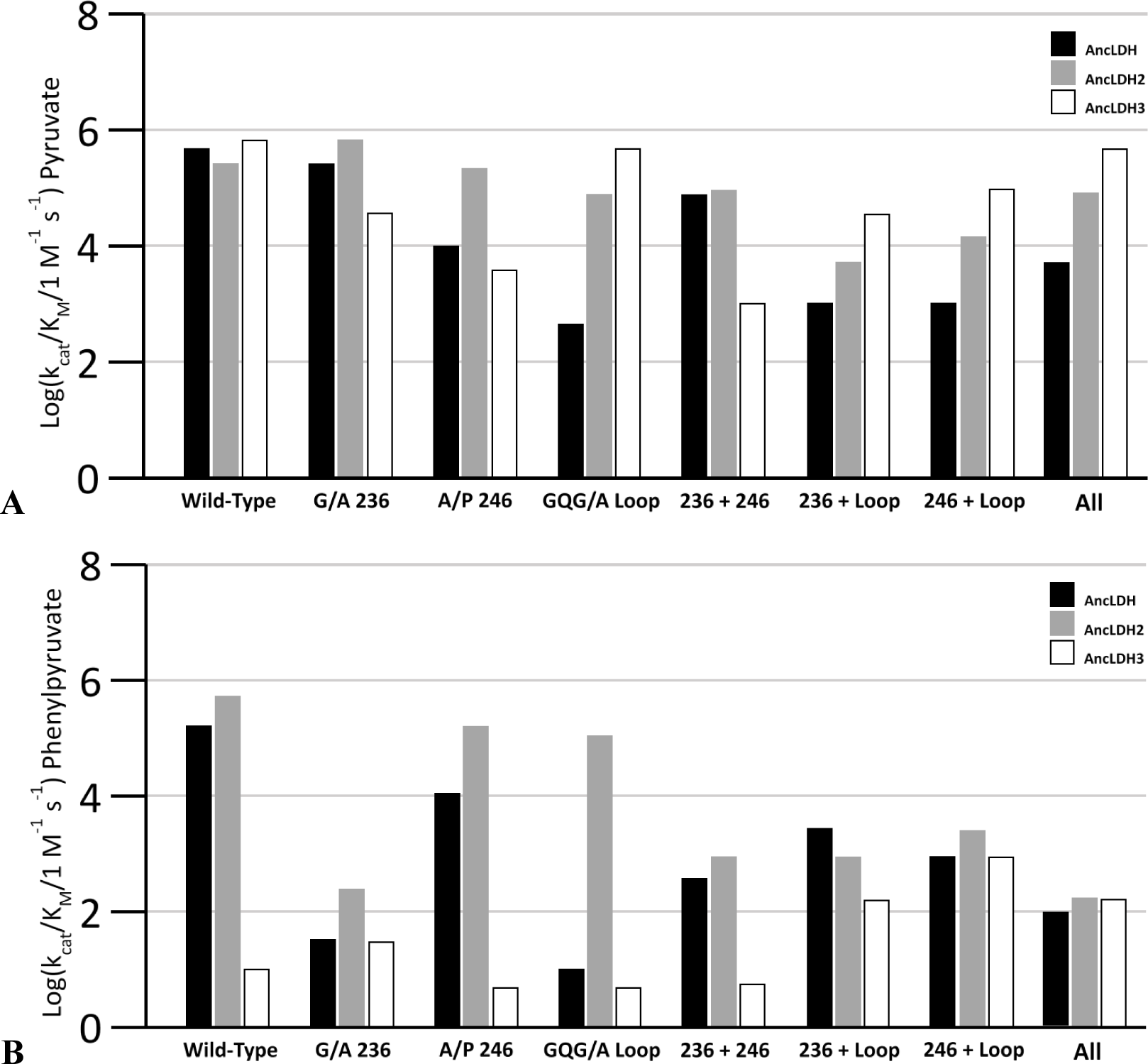
Kinetics of mutant ancestor enzymes **(A)** Pyruvate and **(B)** phenylpyruvate substrate activities are plotted as log(kcat/KM). Each x-axis bin corresponds to the position(s) mutated. ‘All’ refers to the construct that includes both point mutations and the loop mutation. Enzymes are labeled by color.

### Combining Mutations and Apparent Protein Epistasis

AncLDH and AncLDH2 have identical active sites and share 93% sequence identity. The evolutionarily relevant mutations have very different effects in the two ancestors, with changes in AncLDH far more deleterious. It follows that residues within the 7% sequence difference act as permissive mutations by making AncLDH2 more tolerant of active site changes. For example, replacing the GQG motif with an alanine in AncLDH causes a 1000-fold decrease in pyruvate activity and an even greater decrease in activity towards a larger substrate. Doing the equivalent substitution in AncLDH2 has essentially no effect on either substrate, a clear case of protein epistasis (Figure 7, Supp. Table 5 and 6).

Many of the mutation combinations exhibit non-intuitive interactions, especially when using the larger substrate (Figure 7b). In AncLDH2, the G236A swap alone accounts for a ∼3500-fold reduction in phenylpyruvate activity, turnover nearly as slow as with AncLDH3. Coupling G236A with either A246P or the opposing loop mutation, neither having a large impact on their own, reduces the effect to only a few hundred-fold loss in activity (350 and 600, respectively). The equivalent swaps behave similarly in AncLDH3, with each single mutation having amplified (or mitigated) effects when combined that are non-additive.

AncLDH3 was able to recover wild-type pyruvate activity once the entire active site was converted to that of AncLDH2 though curiously some mutant pairs have greatly diminished activity (Figure 7a). Partial rescues in primary substrate activity must be due to epistatic interactions between position 236, 246, and the opposing loop. Interestingly, the AncLDH3 triple mutant only accounted for 100-fold greater phenylpyruvate activity, and so the remaining near-four orders of magnitude difference between the mutant and AncLDH2 must be due to non-active site mutations. Likewise, AncLDH2 cannot be made as specific as AncLDH3 or the modern *Plasmodium* enzyme with active site mutations alone.

## Discussion

### Current Explanations of LDH Specificity Are Incomplete

Epistasis can dictate a protein’s evolutionary trajectory, causing a potentially beneficial mutation in one protein to be harmful in a close relative^2^. LDHs and MDHs were long thought to recognize their substrate solely by using residue 102 on their active site loop. The fact that some canonical LDHs can swap specficities^12, 19^, while others cannot^20-21^, is evidence of epistatic interactions from unknown non-active site residues that help dictate specificity. In recent years, studies have shown that even the classic L/MDH dogma of Q102 vs R102 is not universally correct by characterizing LDHs that use W107f^23^ or L102^14^ as their primary specificity residue.

Apicomplexan LDHs use W107f which was previously thought to hydrophobically pack against the methyl group of pyruvate but sterically block the larger oxaloacetate as a substrate^23^. We have shown here that steric blocking is not a large contribution in the lack of oxaloacetate turnover. Many apicomplexan LDHs can turn over substrates much larger than pyruvate meaning the inability to turnover oxaloacetate is likely only due to charge (Figures 3 and 5). W107f lacks a formal positive charge, which would make oxaloacetate binding disfavored, but the positively charged lysine at position 102 appears positioned such that it could function as a potential charge balancer. The fact that apicomplexan LDHs have poor oxaloacetate despite K102 has been studied but is still unexplained^10, 25, 35-38^.

It is possible that the five amino acid insertion responsible for pyruvate preference displaces position 102 too far from the active site, making K102 unable to balance a negatively charged substrate. However, in Boucher *et al*, a bifunctional ancestral apicomplexan L/MDH required only a single mutation (K102R). Crystal structures of the mutant construct are in the pyruvate-bound conformation, with W107f in the active site and R102 still far from the substrate binding pocket, which should make oxaloacetate unable to fit into the available space. The group’s hypothesis was that the loop could flip either specificity residue into the active site as needed, which is still being tested.

### Plasmodium LDHs Have Atypically High Specificity

LDH_PLFA is unlike its ancestral counterpart because it does not become bifunctional with the K102R mutation, though epistatic ‘locking in’ of function is not an uncommon occurrence^1-2, 14, 23, 48^. LDH_PLFA is locked in to pyruvate preference, has unusually high activity, and is exceptionally specific when compared to close relatives (Figures 3 and 5). Reasons for LDH_PLFA’s remarkable behavior were hypothesized but had not been empirically explored before now^10^. *Plasmodium* LDHs have several mutations that distinguish them from the rest of Apicomplexa and alter the shape of their active sites.

A unique deletion in an active site loop strongly correlates with the inability to use large substrate analogs for turnover (Figure 5 and 6). The two residues 236 and 246 are also unique to *Plasmodium* LDHs and within 4 Å of bound substrate in the closed conformation of the LDH_PLFA (Figure 6). We hypothesize that G236 and A246 which are conserved in other Apicomplexa are more helpful for local flexibility than the *Plasmodium* equivalents A236 and P246. Our data indicates that by evolving these attributes, particularly A236, *Plasmodium* LDHs gained an altered and novel mechanism of substrate recognition, one that includes a strong steric sensor and is absent from other LDHs in the phylum (Figure 5). *Plasmodium* LDHs are more active as well as more specific and it is possible that there is some inherent trade-off between efficiency and promiscuity.

### Epistatic interactions Shape Evolutionary Trajectories

There are a few examples of protein epistasis in our results, both inside and outside the active site. The most obvious is the phenotypic differences between AncLDH mutants and AncLDH2 mutants. The proteins share 93% sequence identity and the differences are exclusively outside of the active site. Mutating the opposing loop motif, however, has drastically different effects: AncLDH was mostly inactivated while AncLDH2 was largely unaffected (Figure 7). The two wild-type enzymes behave similarly meaning some combination within the 7% difference (twenty-two residues) allowed the ancestral LDH to accept the historic opposing loop changes as it evolved towards AncLDH3. The mutation G236A, on the other hand, causes an identical change in phenotype for both ancestors and is the largest source of steric discrimination without affecting pyruvate activity (Figure 7).

The reverse mutation does not confer higher promiscuity on AncLDH3. Instead, A236G has little effect on the enzyme. No combination of active site mutation brings AncLDH3 to ancestral levels promiscuity (Figure 7). The dozens of residues that separate AncLDH3 from the other two ancestral constructs constrain specificity despite all being distant from the active site. Only a few AncLDH3 mutants were crippled with respect to pyruvate activity while changes in in LDH_PLFA were highly detrimental. While not generally uncommon, the behavior surprised us with LDH_PLFA considering how adaptable the enzyme’s primary specificity loop is to mutation^23, 34^.

Numerous epistatic interactions make it hard to identify route the ancestor LDH must have taken during its evolution to arrive at the modern-day sequence. We know that an increase in steric specificity evolved within the *Plasmodium* lineage and that a change should minimally affect the enzyme’s primary role in glycolysis. Given the data, some mutants are more efficient LDHs than others by orders of magnitude implying some evolutionary paths are much less desirable than others. For instance, if AncLDH2’s active site is untouched while the other 90 residues are changed, pyruvate activity is unaffected (Figure 7). There is then only a single active site alteration left available that keeps pyruvate activity at WT levels: G236A. Alternatively, the AncLDH2 active site can mutate through multiple paths with pyruvate activity staying within 10-fold (Figure 7). Some unknown combination of remaining residues increases pyruvate activity and specificity up to AncLDH3 levels.

### Specificity Mechanisms, Conservation, and Inhibitor Design

*P. falciparum* LDH is an active small-molecule drug target because it is essential for the parasite and has an unusual active site architecture^27, 33, 35, 39-45^. The chief difference between *Plasmodium*-like LDHs and host LDH is the specificity loop, its insertion strongly differentiating it from canonical LDHs and making it a target for selective antibody inhibitors as well^49^. Efforts to selectively target the apicomplexan specificity loop itself may be ineffective, however, due to the lack of functional constraint. Recent work has outlined that the evolution of the loop insertion was largely noncontingent, with its unique length and sequence of little importance, implying a large potential for the development of resistance towards loop-targeting drugs^34^.

High shared sequence identity within apicomplexan LDHs has led to the belief that successful small-molecule inhibitors selectively targeting the active site (but not necessarily the loop) would likely be effective at treating other apicomplexan-caused diseases^33, 35-36, 38-39, 43-44^. However, the evolution of a unique substrate recognition mechanism would mean inhibitor interactions studied in a *Plasmodium* LDH model may not be conserved across Apicomplexa. Just as likely, inhibitors robust towards Apicomplexa such as *Toxoplasma* and *Eimeria* may be rendered ineffective for *Plasmodium* due to different specificity mechanisms.

*P. falciparum* LDH’s strong specificity makes rational inhibitor design difficult without further studying its source. The straightforward approach reported here revealed that the enzyme is unable to reduce large substrate analogs using NADH but gave little insight into effects on binding potential. AncLDH3 has a K_M_ value for phenylpyruvate 10-fold weaker than AncLDH2 but a k_cat_ 3600-fold slower (Figure 7). If the rate limiting step in the ancestors is also closure of the specificity loop, large substrates are possibly rejected due to the inability for the loop close and occlude water proficiently. Active site substitutions in AncLDH2 that lower activity towards large substrates without affecting pyruvate rates accomplish it primarily through reducing k_cat_ while K_M_ is relatively unaffected.

Slow loop closure could allow substrate to dissociate more easily before catalysis, causing a sufficiently large increase in k_off_ that the observed K_M_ remains the same^50^. This suggests that it will be highly difficult to find effective inhibitors or other substrate analogs that selectively target the unique *Plasmodium* LDH active site. The data presented here is insufficient to elucidate true binding affinity, however. Whether a change in k_off_ or k_on_ was the key factor for *Plasmodium* when evolving such strong substrate preference is unknown and those hypotheses must be tested in the future.

## Methods

### Phylogenetics

Protein sequences used for phylogenetic tree construction were obtained using either the RefSeq database^51^ with the BLASTP search algorithm^52^ using the chosen query sequences. Four query sequences were used to ensure sufficient coverage for the superfamily: UniProtIDs P11708, Q76NM3, C6KT25, and Q73G44. Redundant sequences, constructs, and PDB sequences were removed. Sequences were curated based on length and sequence identity in order to reduce computation time. The dataset was trimmed down to 60% sequence identity with apicomplexan sequences reintroduced to ensure full coverage of the nodes of interest. The final dataset had 277 taxa. A sequence alignment was generated with MUSCLE and a ML tree was inferred using PhyML^53^ using the LG substitution matrix^54^ and estimating the gamma parameter using 12 categories.

### Ancestral Sequence Reconstruction

Sequences at internal branch points in the phylogenies were reconstructed using the codeml function in the PAML program suite^55^. Posterior amino acid probabilities were calculated using the LG substitution matrix^54^ given the tree generated by PhyML^53^. The reconstructions estimated background frequencies of amino acids from the alignment. To assist with proper gene expression, the N- and C-termini were modified manually to match the closest modern sequence, which was determined by branch length.

### Plasmid Construction and Mutagenesis

For all proteins, codon optimized genes were synthesized by GenScript (Piscataway, NJ) and sub-cloned into pET-24a, between the NdeI and XhoI restriction sites, without the N-terminal T7-tag but using the C-terminal 6xHistidine-tag. All point mutations were made using the QuikChange Lightning kit from Agilent (Santa Clara, CA) and all relevant mutagenic primers were synthesized by IDT (Coralville, IA). Larger mutations (Indels >1 amino acid) were performed by GenScript directly using the previously synthesized construct as a template. Sequences were confirmed by Sanger Sequencing at Genewiz (Cambridge, Massachusetts). Sequence numbering for all mutations is taken directly from the sequence alignment.

### Protein Expression and Purification

All materials were purchased from Sigma-Aldrich (St. Louis, MO) unless otherwise stated. Plasmids were transformed into BL21-DE3 (pLysS) *E. coli* cells (Invitrogen, Grand Island, NY) for expression. Cells were grown in a shaker-incubator at 37 °C, 225 RPM agitation in 2xYT media supplemented with 30 mM KH_2_PO_4_, pH 7.8, 0.1% (w/v) glucose and cell growth was monitored at OD_600_. Once an OD_600_ of 0.5-0.8 was reached, protein expression was induced by adding 0.5 mM isopropyl β-d-1-thiogalactopyranoside to each culture and they were further incubated for 4 hours at 37 °C with 225 RPM agitation. Cells were pelleted by centrifugation and stored at −80 °C.

Pellets were thawed on ice and resuspended in 20 mL of HisTrap Binding Buffer (50 mM NaH_2_PO_4_, pH 7.4, 300 mM NaCl, 10 mM imidazole) and 2 µL Pierce Universal Nuclease (Thermo Scientific, Rockford, IL). Once resuspended, lysate was sonicated on ice at 35% amplitude in pulses of 30 seconds ON and 20 seconds OFF for 2 minutes. Insoluble cell debris was separated by centrifugation at 18,000*xg* for 20 minutes. The supernatant was syringe-filtered to 0.22 µm and then purified using a 5 mL HisTrap FF nickel affinity column (GE Healthcare, Piscataway, NJ). Fractions were eluted with an imidazole buffer (50 mM NaH_2_PO_4_, pH 7.4, 300 mM NaCl, 500 mM imidazole) on an AKTA Prime Plus (GE Healthcare, Piscataway, NJ). Fractions were analyzed by SDS-PAGE, pooled, and concentrated using Amicon Ultracel-10K centrifugation filter units (Millipore, Billerica, MA). Proteins were buffer-exchange into 50 mM Tris, pH 7.4, 100 mM NaCl, 0.1 mM EDTA, 0.01% azide by a PD-10 column (GE Healthcare, Piscataway, NJ). Enzyme concentrations were determined by the sample’s 280 nm absorbance, using extinction coefficients and molecular weights calculated by ExPASy’s ProtParam tool.

### **X-** ray Crystallography

Conditions were optimized based on promising hits identified from screens with Crystal Screen and Crystal Screen 2 (Hampton Research, Aliso Viejo, CA). Proteins were crystallized by hanging-drop vapor diffusion (2 µl protein stock and 2 µl well solution) in VDX greased plates and siliconized cover slides (Hampton) and equilibrated against 1 mL of the well solution at room temperature. Crystals were cryo-protected by being soaked in a 15% (w/v) dextrose solution for 3 minutes, transferred to a 30% (w/v) dextrose solution, and then quickly flash-frozen in liquid N_2_.

Diffraction datasets were collected at the SIBYLS beamline (Lawrence Berkeley National Laboratory, Berkeley, CA). All data was collected at 100 K, indexed, integrated, and called with XDS/XSCALE^56^, and all reflections with a CC(1/2) above 10% were used. Molecular replacement and refinement using these reflections were done with the PHENIX software suite^57^. Modeling was performed using Coot^58^ and model quality was validated with MolProbity^59-60^. Superpositions were generated using THESEUS^61^ and images were rendered with PyMOL^62^. In solving the structure 6CT6, the active site was not fully closed, and accordingly has a disordered loop with high B-factors. The electron density in this region is poor and insufficient to support modeling the structure of the entire loop. Consequently, a few residues are missing from the solved structures.

### Steady-State Kinetics Assays

Enzymes were assayed at 25 °C by recording the change in absorbance at 340 nm for 300 seconds on a Cary 100 Bio (Agilent, Santa Clara, CA) in 50 mM Tris, pH 7.5, 50 mM KCl^10, 14, 23, 34^. Enzyme concentrations varied as needed and ranged from 1 nM to 10 μM. The NADH concentration was a constant 200 μM, except where noted, well above saturating for most MDHs^63^ and LDHs^64^, while substrate concentrations were varied. All experiments monitored the oxidation of NADH unless stated otherwise. Data was fit together with Kaleidegraph as a chi-squared estimate of either the Michaelis-Menten equation or a Substrate Inhibition model:

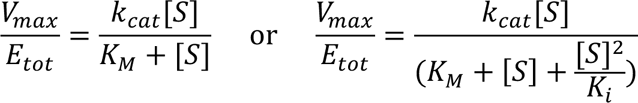

The error for each parameter is reported as the standard error of the mean (SEM) from triplicate measurements as estimated by the fitting software.

Oxaloacetate spontaneously decarboxylates to pyruvate at 25 °C under neutral aqueous conditions at a significant rate^65^. Oxaloacetate preparations are known to contain pyruvate contamination (a few percent in purchases from Sigma-Aldrich, dependent on the batch) which must be kept in mind when assaying promiscuous L/MDHs. All oxaloacetate stocks were made fresh directly before each experiment and stored on ice throughout. Enzymes that behave as ‘poor’ LDHs are relatively unaffected by the contaminant. Enzymes that have high pyruvate activity or are especially promiscuous can appear to have high artifactual oxaloacetate activity that is due to pyruvate turnover^10, 12, 23, 66^. Therefore, oxaloacetate activity for primary LDHs was assayed at high enzyme concentrations (micromolar amounts) which results in a non-linear biphasic decrease in A_340_. The initial burst corresponds to pyruvate contamination quickly being turned over by the enzyme and the slow, second linear portion was used as the apparent oxaloacetate reduction^12, 66^.

## Supplemental Information

**Supp. Table 1.** Alternative Substrate Kinetic Parameters

| Enzyme | LDH_PLFA |  |  | LDH_EIMA |  |  | Anc_LDH |  |  |
| --- | --- | --- | --- | --- | --- | --- | --- | --- | --- |
| Substrate | $k_{cat}$<br>(s<sup>-1</sup>) | $K_M$<br>(μM<sup>-1</sup>) | $k_{cat}/K_M$<br>(M<sup>-1</sup> s<sup>-1</sup>) | $k_{cat}$<br>(s<sup>-1</sup>) | $K_M$<br>(μM<sup>-1</sup>) | $k_{cat}/K_M$<br>(M<sup>-1</sup> s<sup>-1</sup>) | $k_{cat}$<br>(s<sup>-1</sup>) | $K_M$<br>(μM<sup>-1</sup>) | $k_{cat}/K_M$<br>(M<sup>-1</sup> s<sup>-1</sup>) |
| Pyruvate | 91<br>±3 | 72<br>±9 | 1.3x10<sup>6</sup> | 7.4<br>±0.2 | 57<br>±4 | 1.3x10<sup>5</sup> | 45<br>±3 | 100<br>±30 | 4.2x10<sup>5</sup> |
| Phenylpyruvate | 0.03<br>±0.01 | 1500<br>±200 | 2.0x10<sup>1</sup> | 16<br>±0.4 | 560<br>±60 | 2.8x10<sup>4</sup> | 52<br>±2 | 440<br>±80 | 1.2x10<sup>5</sup> |
| α-ketovalerate | 0.10<br>±0.01 | 2400<br>±300 | 5.7x10<sup>1</sup> | 3.1<br>±0.1 | 3200<br>±400 | 9.7x10<sup>2</sup> | 9.5<br>±0.4 | 3000<br>±500 | 3.1x10<sup>3</sup> |
| Isocaproate | ND | ND | ND | 0.42<br>±0.01 | 2700<br>±400 | 1.6x10<sup>2</sup> | 2.0<br>±0.1 | 2500<br>±500 | 8.0x10<sup>2</sup> |
| Oxaloacetate | ND | ND | 1.0x10<sup>1</sup> | ND | ND | 3.8x10<sup>1</sup> | 0.37<br>±0.01 | 6500<br>±500 | 5.6x10<sup>1</sup> |

**Supp. Table 2.**
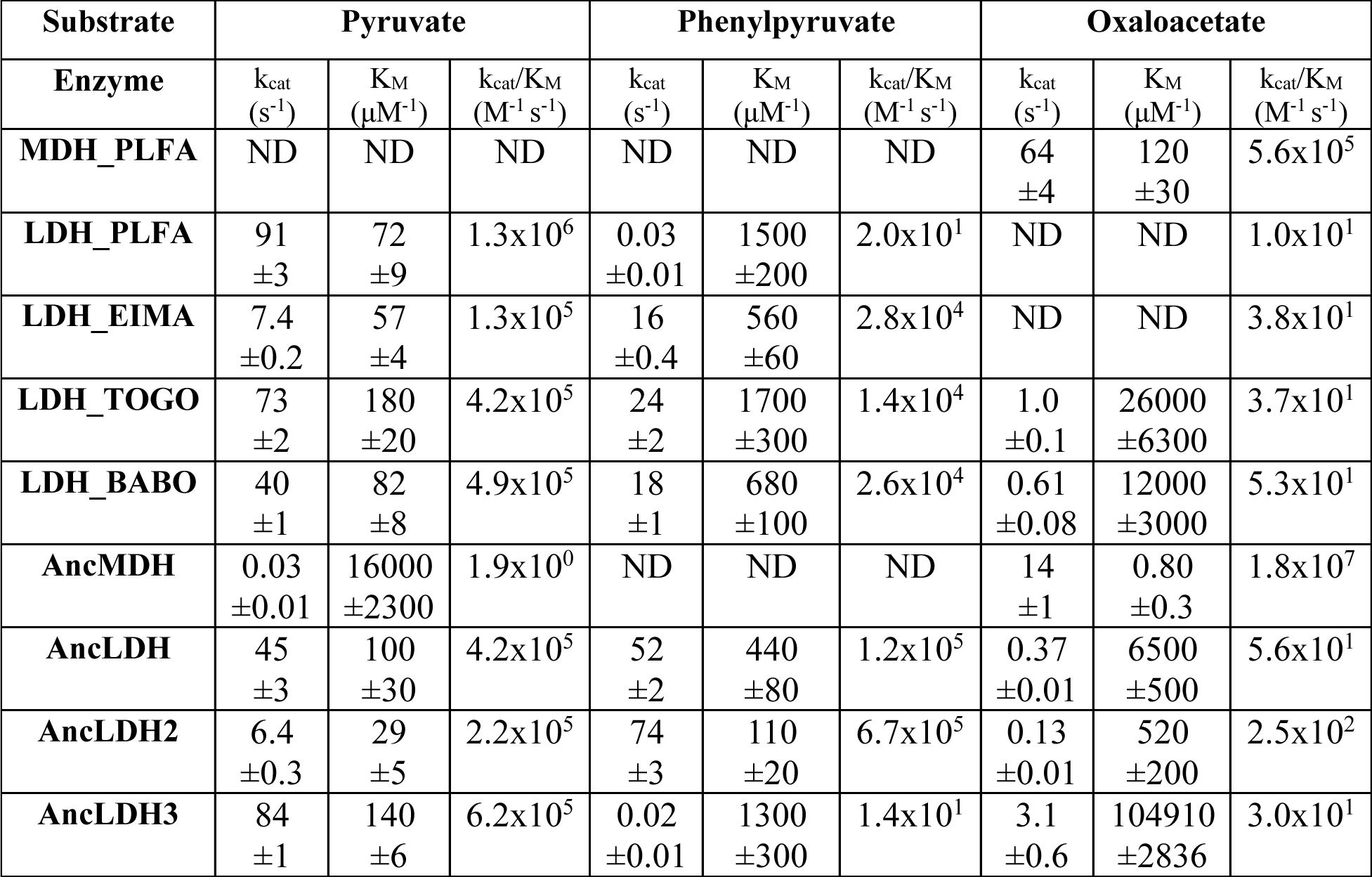
Steric Specificity Kinetics in Apicomplexan LDHs

| Substrate | Pyruvate |  |  | Phenylpyruvate |  |  | Oxaloacetate |  |  |
| --- | --- | --- | --- | --- | --- | --- | --- | --- | --- |
| Enzyme | $k_{cat}$<br>(s<sup>-1</sup>) | $K_M$<br>(μM<sup>-1</sup>) | $k_{cat}/K_M$<br>(M<sup>-1</sup> s<sup>-1</sup>) | $k_{cat}$<br>(s<sup>-1</sup>) | $K_M$<br>(μM<sup>-1</sup>) | $k_{cat}/K_M$<br>(M<sup>-1</sup> s<sup>-1</sup>) | $k_{cat}$<br>(s<sup>-1</sup>) | $K_M$<br>(μM<sup>-1</sup>) | $k_{cat}/K_M$<br>(M<sup>-1</sup> s<sup>-1</sup>) |
| MDH_PLFA | ND | ND | ND | ND | ND | ND | 64<br>±4 | 120<br>±30 | 5.6x10<sup>5</sup> |
| LDH_PLFA | 91<br>±3 | 72<br>±9 | 1.3x10<sup>6</sup> | 0.03<br>±0.01 | 1500<br>±200 | 2.0x10<sup>1</sup> | ND | ND | 1.0x10<sup>1</sup> |
| LDH_EIMA | 7.4<br>±0.2 | 57<br>±4 | 1.3x10<sup>5</sup> | 16<br>±0.4 | 560<br>±60 | 2.8x10<sup>4</sup> | ND | ND | 3.8x10<sup>1</sup> |
| LDH_TOGO | 73<br>±2 | 180<br>±20 | 4.2x10<sup>5</sup> | 24<br>±2 | 1700<br>±300 | 1.4x10<sup>4</sup> | 1.0<br>±0.1 | 26000<br>±6300 | 3.7x10<sup>1</sup> |
| LDH_BABO | 40<br>±1 | 82<br>±8 | 4.9x10<sup>5</sup> | 18<br>±1 | 680<br>±100 | 2.6x10<sup>4</sup> | 0.61<br>±0.08 | 12000<br>±3000 | 5.3x10<sup>1</sup> |
| AncMDH | 0.03<br>±0.01 | 16000<br>±2300 | 1.9x10<sup>0</sup> | ND | ND | ND | 14<br>±1 | 0.80<br>±0.3 | 1.8x10<sup>7</sup> |
| AncLDH | 45<br>±3 | 100<br>±30 | 4.2x10<sup>5</sup> | 52<br>±2 | 440<br>±80 | 1.2x10<sup>5</sup> | 0.37<br>±0.01 | 6500<br>±500 | 5.6x10<sup>1</sup> |
| AncLDH2 | 6.4<br>±0.3 | 29<br>±5 | 2.2x10<sup>5</sup> | 74<br>±3 | 110<br>±20 | 6.7x10<sup>5</sup> | 0.13<br>±0.01 | 520<br>±200 | 2.5x10<sup>2</sup> |
| AncLDH3 | 84<br>±1 | 140<br>±6 | 6.2x10<sup>5</sup> | 0.02<br>±0.01 | 1300<br>±300 | 1.4x10<sup>1</sup> | 3.1<br>±0.6 | 104910<br>±2836 | 3.0x10<sup>1</sup> |

**Supp. Figure 1.**
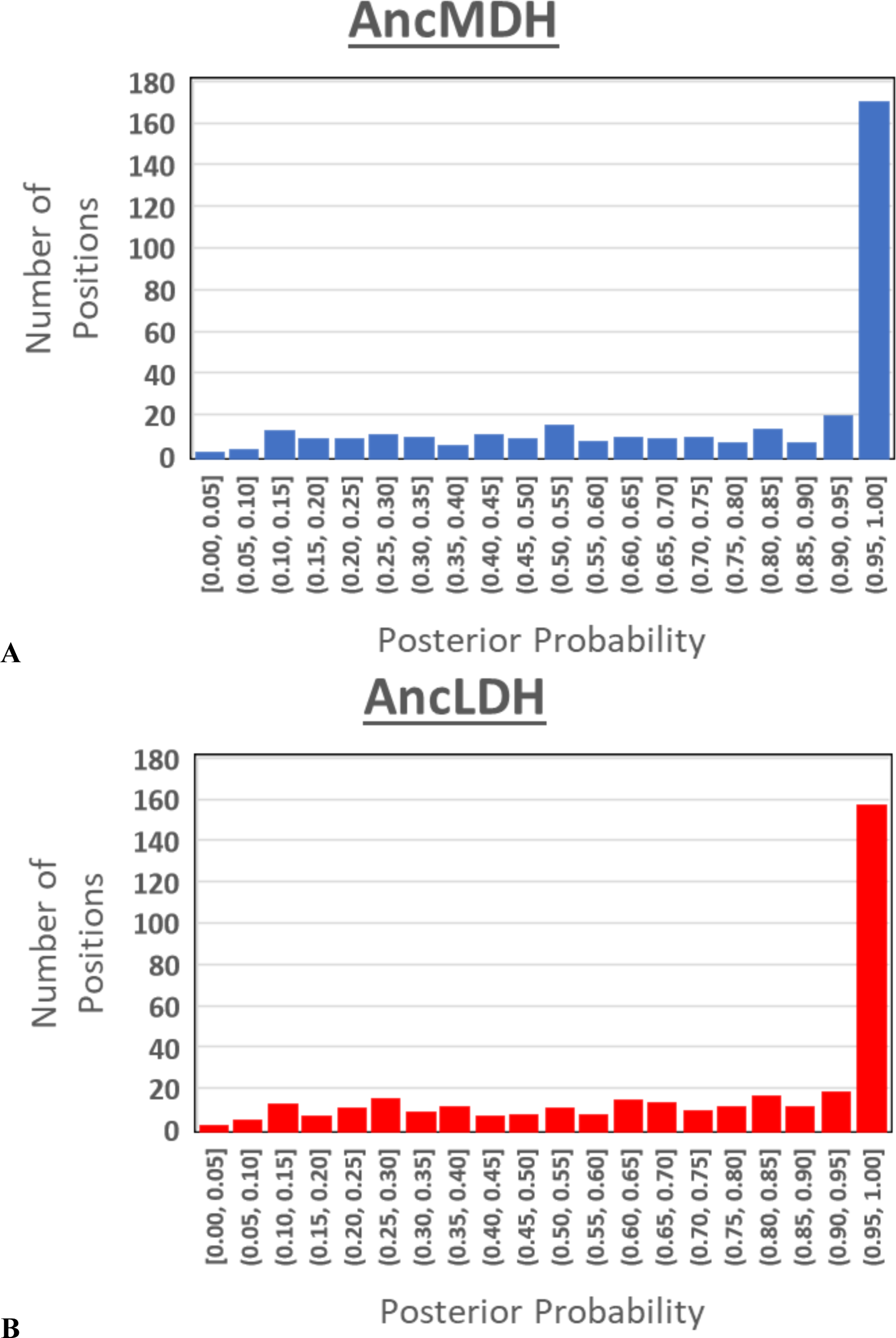
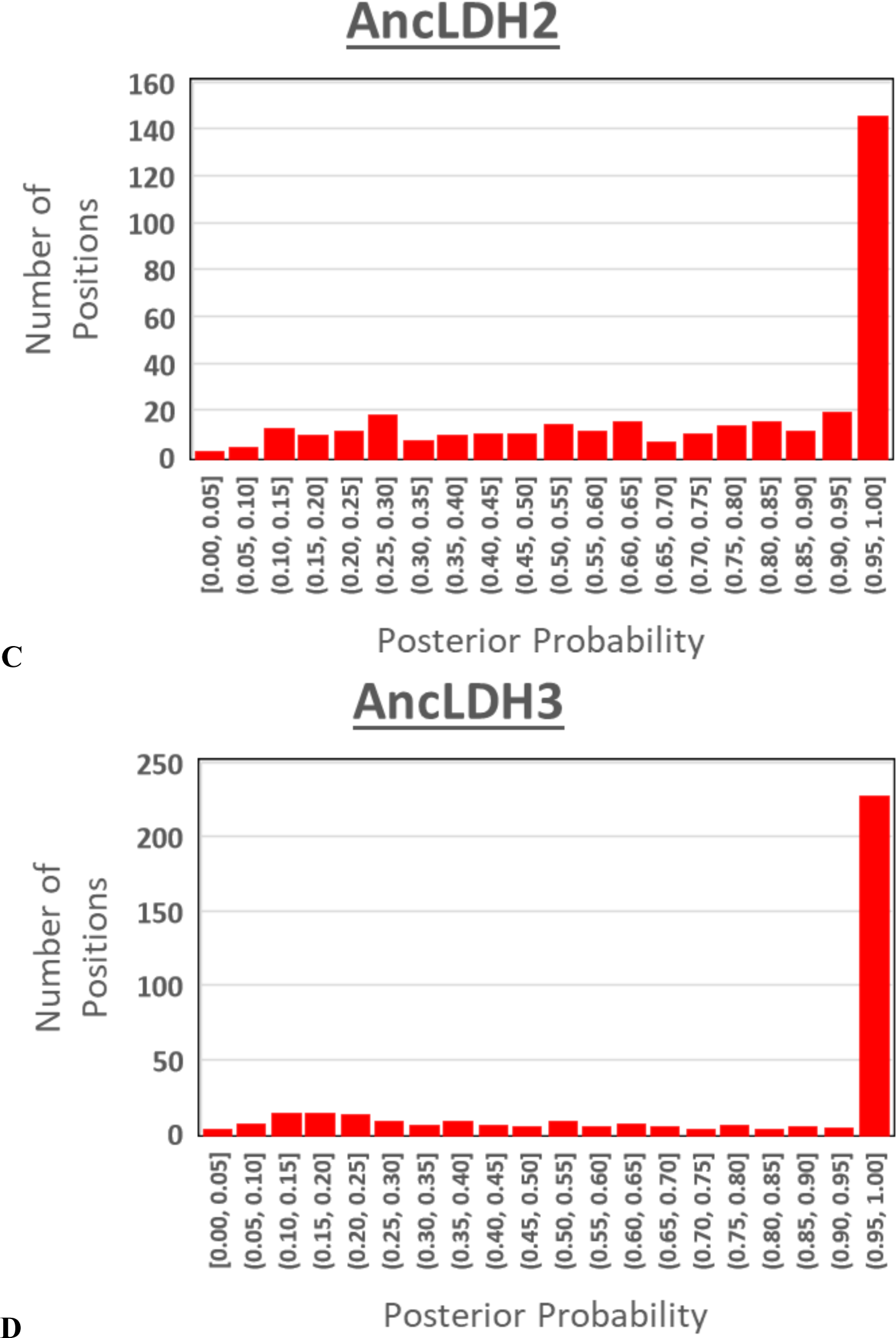
Posterior probabilities histograms. Each amino acid site in the ancestral reconstructions are binned according to the posterior probability of the predicted residue. **(A)** AncMDH, **(B)** AncLDH, **(C)** AncLDH2, and **(D)** AncLDH3 are colored by primary substrate preference.

**Supp. Table 3.** Crystallographic Statistics for LDH_EIMA

| **Data Collection** | LDH_EIMA (6CT6) |
| --- | --- |
| Space group | P 31 2 1 |
| Cell dimensions |  |
| *a, b, c* (Å) | 83.4 83.4 227.1 |
| $\alpha, \beta, \gamma$ (°) | 90 90 120 |
| Resolution (high res. bin) (Å) | 44.64 - 1.705 (1.729 - 1.705) |
| I/σI (high res. bin) | 6.43 (0.34) |
| R<sub>meas</sub> | 0.134 (3.66) |
| CC(1/2) | 100 (66.6) |
| Resolution at CC(1/2) = 0.50 (Å) | 1.05 |
| Completeness (%) | 88.68 (98.5) |
| Total observations | 170863 |
| Unique observations | 100328 (9532) |
| Redundancy | 1.84 |
| Wilson B-factor | 30.76 |
| Collected at beamline | ALS 12.3.1 |
| **Model Refinement** |  |
| Resolution (Å) | 45.7 – 1.05 |
| R<sub>work</sub>/R<sub>free</sub> | 0.2016 / 0.2183 |
| R<sub>work</sub>/R<sub>free</sub> at CC(1/2) = 0.5 res. cutoff | 0.2016 / 0.2183 |
| Mol. Per ASU (protein + ligand) | 2 |
| Number of atoms | 5438 |
| protein | 4958 |
| ligand/ions | 74 |
| waters | 406 |
| Average B-factor | 14.0 |
| r.m.s. deviations |  |
| Bond lengths (Å) | 0.009 |
| Bond angles (Å) | 0.79 |
| Ramachandran favored (%) | 97.04 |
| Ramachandran outliers (%) | 0.16 |
| All-atom clashscore | 0.2 |

**Supp. Table 4.** Sequence Identity of Modern and Ancestor Proteins

| Enzyme | MDH<br>PLFA | LDH<br>PLFA | LDH<br>EIMA | LDH<br>TOGO | LDH<br>BABO | Anc<br>MDH | Anc<br>LDH | Anc<br>LDH2 | Anc<br>LDH3 |
| --- | --- | --- | --- | --- | --- | --- | --- | --- | --- |
| MDH_PLFA | - | 40% | 42% | 40% | 38% | 51% | 48% | 46% | 40% |
| LDH_PLFA | 40% | - | 52% | 49% | 51% | 65% | 63% | 66% | 91% |
| LDH_EIMA | 42% | 52% | - | 59% | 50% | 56% | 65% | 61% | 53% |
| LDH_TOGO | 40% | 49% | 59% | - | 48% | 55% | 64% | 55% | 50% |
| LDH_BABO | 38% | 51% | 50% | 48% | - | 55% | 62% | 67% | 51% |
| AncMDH | 51% | 65% | 56% | 55% | 55% | - | 82% | 78% | 57% |
| AncLDH | 48% | 63% | 65% | 64% | 62% | 82% | - | 93% | 66% |
| AncLDH2 | 46% | 66% | 61% | 55% | 67% | 78% | 93% | - | 70% |
| AncLDH3 | 40% | 91% | 53% | 50% | 51% | 57% | 66% | 70% | - |

**Supp. Table 5.** Single Mutant Ancestral Apicomplexan LDH Kinetics

| Substrate | Pyruvate |  |  | Phenylpyruvate |  |  | Oxaloacetate |  |  |
| --- | --- | --- | --- | --- | --- | --- | --- | --- | --- |
| Enzyme | k<sub>cat</sub><br>(s<sup>-1</sup>) | K<sub>M</sub><br>(μM<sup>-1</sup>) | k<sub>cat</sub>/K<sub>M</sub><br>(M<sup>-1</sup> s<sup>-1</sup>) | k<sub>cat</sub><br>(s<sup>-1</sup>) | K<sub>M</sub><br>(μM<sup>-1</sup>) | k<sub>cat</sub>/K<sub>M</sub><br>(M<sup>-1</sup> s<sup>-1</sup>) | k<sub>cat</sub><br>(s<sup>-1</sup>) | K<sub>M</sub><br>(μM<sup>-1</sup>) | k<sub>cat</sub>/K<sub>M</sub><br>(M<sup>-1</sup> s<sup>-1</sup>) |
| AncLDH<br>G236A | 100<br>±4 | 470<br>±70 | 2.3x10<sup>5</sup> | 0.10<br>±0.01 | 3500<br>±600 | 3.5x10<sup>1</sup> | 0.31<br>±0.02 | 7500<br>±1000 | 4.2x10<sup>1</sup> |
| AncLDH2<br>G236A | 37<br>±1 | 54<br>±8 | 6.7x10<sup>5</sup> | 0.30<br>±0.01 | 220<br>±200 | 1.8x10<sup>2</sup> | 0.20<br>±0.01 | 5400<br>±700 | 3.9x10<sup>1</sup> |
| AncLDH3<br>A236G | 23<br>±1 | 600<br>±100 | 3.8x10<sup>4</sup> | 0.09<br>±0.01 | 2100<br>±700 | 4.1 x10<sup>1</sup> | ND | ND | 4.9x10<sup>1</sup> |
| AncLDH<br>A246P | 15<br>±1 | 2400<br>±400 | 6.0x10<sup>3</sup> | 20<br>±1 | 1700<br>±200 | 1.2x10<sup>4</sup> | 0.68<br>±0.02 | 12000<br>±2000 | 5.6x10<sup>1</sup> |
| AncLDH2<br>A246P | 15<br>±0.5 | 230<br>±20 | 6.2x10<sup>4</sup> | 48<br>±2 | 500<br>±60 | 9.6x10<sup>4</sup> | 0.20<br>±0.01 | 2600<br>±400 | 6.7x10<sup>1</sup> |
| AncLDH3<br>P246A | 7.3<br>±0.3 | 1600<br>±200 | 4.6x10<sup>3</sup> | 0.04<br>±0.01 | 6600<br>±3000 | 6.0x10<sup>0</sup> | 5.5<br>±3 | 200000<br>±100000 | 2.8x10<sup>1</sup> |
| AncLDH<br>Deletion | 14<br>±1 | 25000<br>±4000 | 5.7x10<sup>2</sup> | 0.10<br>±0.01 | 2300<br>±300 | 1.7x10<sup>1</sup> | 0.31<br>±0.02 | 10000<br>±2000 | 3.0x10<sup>1</sup> |
| AncLDH2<br>Deletion | 39<br>±1 | 730<br>±70 | 5.3x10<sup>4</sup> | 75<br>±2 | 1300<br>±200 | 5.8x10<sup>4</sup> | 0.50<br>±0.03 | 8900<br>±900 | 5.5x10<sup>1</sup> |
| AncLDH3<br>Insert | 71<br>±2 | 130<br>±10 | 5.4x10<sup>5</sup> | 0.10<br>±0.02 | 15000<br>±6000 | 6.5x10<sup>0</sup> | 5.5<br>±3 | 200000<br>±100000 | 2.8x10<sup>1</sup> |
| AncLDH<br>Combined | 20<br>±1 | 3300<br>±600 | 5.9x10<sup>3</sup> | 0.20<br>±0.01 | 2400<br>±200 | 9.4x10<sup>1</sup> | 1.2<br>±0.2 | 45000<br>±10000 | 2.8x10<sup>1</sup> |
| AncLDH2<br>Combined | 5.1<br>±1 | 130<br>±20 | 3.9x10<sup>4</sup> | 0.60<br>±0.04 | 2100<br>±600 | 2.9x10<sup>2</sup> | 0.05<br>±0.03 | 9600<br>±2000 | 5.2x10<sup>1</sup> |
| AncLDH3<br>Combined | 16<br>±0.4 | 63<br>±5 | 2.5x10<sup>5</sup> | 0.24<br>±0.01 | 1100<br>±100 | 2.1x10<sup>2</sup> | 2.9<br>±0.6 | 96000<br>±30000 | 3.0x10<sup>1</sup> |

**Supp. Table 6.** Double Mutant Ancestral Apicomplexan LDH Kinetics

| Substrate | Pyruvate |  |  | Phenylpyruvate |  |  | Oxaloacetate |  |  |
| --- | --- | --- | --- | --- | --- | --- | --- | --- | --- |
| Enzyme | $k_{cat}$<br>(s<sup>-1</sup>) | $K_M$<br>(μM<sup>-1</sup>) | $k_{cat}/K_M$<br>(M<sup>-1</sup> s<sup>-1</sup>) | $k_{cat}$<br>(s<sup>-1</sup>) | $K_M$<br>(μM<sup>-1</sup>) | $k_{cat}/K_M$<br>(M<sup>-1</sup> s<sup>-1</sup>) | $k_{cat}$<br>(s<sup>-1</sup>) | $K_M$<br>(μM<sup>-1</sup>) | $k_{cat}/K_M$<br>(M<sup>-1</sup> s<sup>-1</sup>) |
| **AncLDH**<br>**G236A/A246P** | 57<br>±1 | 820<br>±80 | 7.0x10<sup>4</sup> | 0.90<br>±0.1 | 1300<br>±300 | 6.4x10<sup>2</sup> | 0.32<br>±0.02 | 5000<br>±800 | 6.4x10<sup>1</sup> |
| **AncLDH2**<br>**G236A/A246P** | 6.3<br>±4 | 83<br>±20 | 7.7x10<sup>4</sup> | 2.1<br>±0.1 | 1700<br>±300 | 2.0x10<sup>3</sup> | 0.10<br>±0.01 | 1800<br>±300 | 7.0x10<sup>1</sup> |
| **AncLDH3**<br>**A236G/P246A** | 4.1<br>±0.2 | 1800<br>±300 | 2.2x10<sup>3</sup> | 0.03<br>±0.01 | 2200<br>±400 | 1.1x10<sup>1</sup> | 3.0<br>±0.8 | 94000<br>±40000 | 3.1x10<sup>1</sup> |
| **AncLDH**<br>**G236A/Del** | 63<br>±10 | 26000<br>±8000 | 2.4x10<sup>3</sup> | 16<br>±2 | 3200<br>±300 | 5.1x10<sup>3</sup> | 0.80<br>±0.1 | 31000<br>±5000 | 2.6x10<sup>1</sup> |
| **AncLDH2**<br>**G236A/Del** | 12<br>±0.03 | 1300<br>±100 | 9.0x10<sup>3</sup> | 1.9<br>±0.1 | 1800<br>±300 | 1.2x10<sup>3</sup> | 0.50<br>±0.01 | 2000<br>±200 | 2.4x10<sup>2</sup> |
| **AncLDH3**<br>**A236G/Ins** | 38<br>±8 | 870<br>±70 | 4.3x10<sup>4</sup> | 0.20<br>±0.01 | 870<br>±200 | 2.3x10<sup>2</sup> | ND | ND | 2.4x10<sup>1</sup> |
| **AncLDH**<br>**A246P/Del** | 15<br>±0.8 | 5400<br>±1000 | 2.8x10<sup>3</sup> | 6.0<br>±0.3 | 3000<br>±500 | 1.9x10<sup>3</sup> | 0.50<br>±0.03 | 25000<br>±3000 | 1.9x10<sup>1</sup> |
| **AncLDH2**<br>**A246P/Del** | 10<br>±0.3 | 460<br>±60 | 2.2x10<sup>4</sup> | 30<br>±6 | 3200<br>±900 | 9.3x10<sup>3</sup> | 0.60<br>±0.1 | 13000<br>±3000 | 4.3x10<sup>1</sup> |
| **AncLDH3**<br>**P246A/Ins** | 22<br>±0.4 | 140<br>±10 | 1.5x10<sup>5</sup> | 0.01<br>±0.01 | 2300<br>±400 | 3.7x10<sup>0</sup> | 1.9<br>±0.3 | 57000<br>±30000 | 3.4x10<sup>1</sup> |

